# Characterizing the Molecular Determinants of Clamp Binding in *B. subtilis*

**DOI:** 10.64898/2026.05.27.728225

**Authors:** Sarah J. Rancic, Katrin M. Klassen, Nicholas Sawyer, Elizabeth S. Thrall

## Abstract

In bacteria, the ring-shaped sliding clamp, DnaN, is an essential component of the replication machinery. The clamp encircles the parental DNA strand during replication and binds DNA polymerases and other replication and repair proteins, helping to tether them at their site of action on the DNA strand. These binding partners interact with the clamp via short pentapeptide or hexapeptide sequences known as clamp-binding motifs (CBMs). Although conserved CBM sequences have been identified across different bacterial species, most studies of clamp binding have been performed in the model gram-negative bacterium *Escherichia coli* and less is known about clamp binding in other bacterial species. In this study, we investigate clamp binding in the model gram-positive bacterium *Bacillus subtilis*. We use fluorescence polarization binding assays to quantify binding of a range of CBM peptides to the clamps of both *E. coli* and *B. subtilis*. We identify similarities in clamp binding between the two species, including similar importance of different amino acids within the conserved pentapeptide motif. However, our results also reveal differences in clamp binding between the two species. Most notably, we find that, although pentapeptide CBMs bind the *E. coli* and *B. subtilis* clamps with similar affinity, hexapeptide CBMs bind an order of magnitude more weakly to the *B. subtilis* clamp. Our results provide new insight into clamp binding in bacteria and point to possible species-specific differences in this essential interaction.

## TEXT

DNA replication is carried out by a multi-protein complex known as the replisome.^1,2^ One replisome component, the sliding clamp, is an essential replication accessory factor across all domains of life.^3,4^ Sliding clamps are ring-shaped proteins that encircle the DNA template, tethering DNA polymerases and other binding partners to their site of action. In bacteria, the clamp is known as DnaN (or β in *E. coli*); it is a heterodimer composed of two identical protomers (Figure 1A). Clamp binding is generally essential for the function of replicative DNA polymerases, and it is also important in DNA repair and damage tolerance pathways.^5^

**Figure 1.**
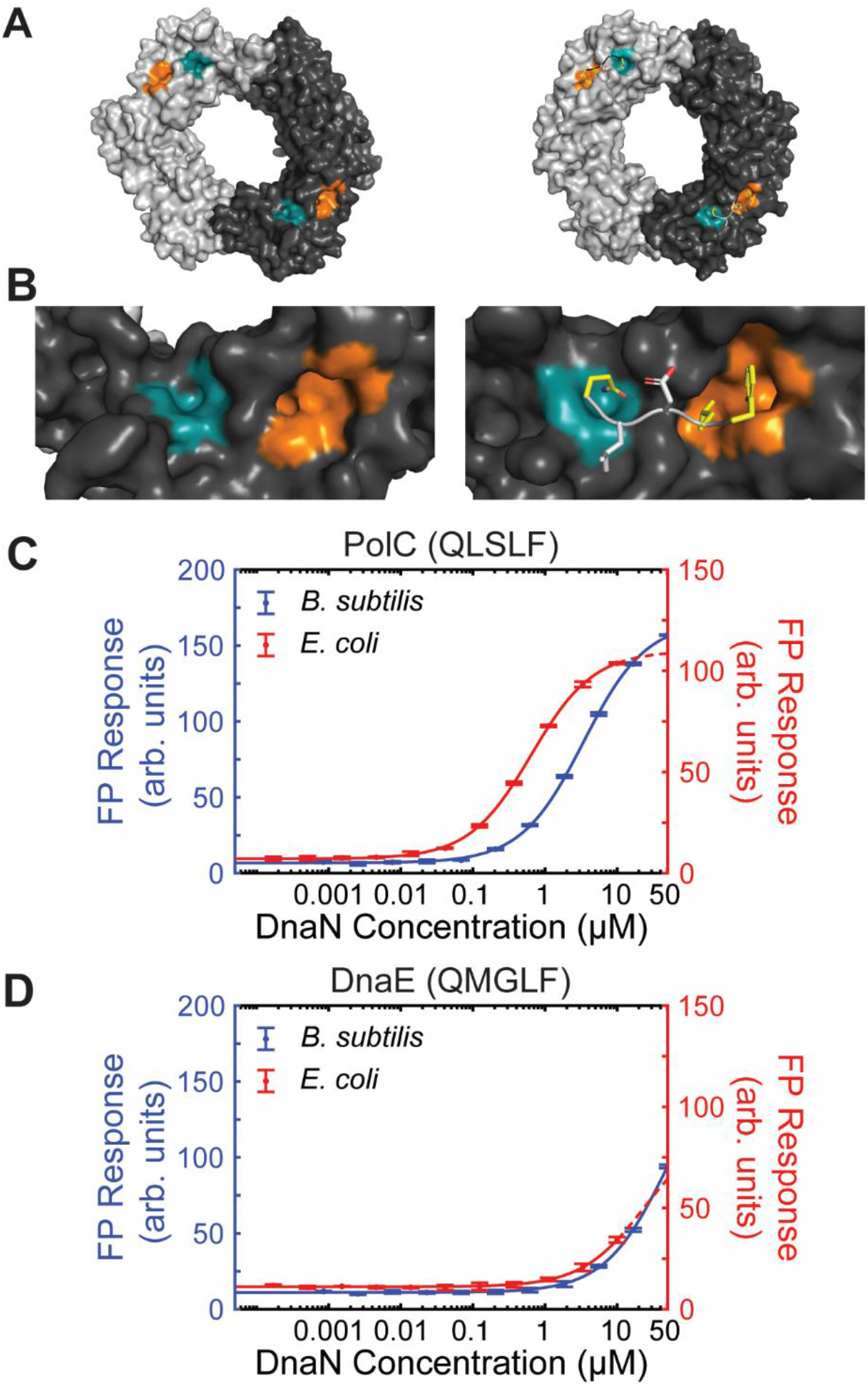
Clamp structures and FP response for DNA polymerase CBM peptides. (A) Crystal structures of (left) *B. subtilis* DnaN with empty binding pocket (PDB: 4TR6) and (right) *E. coli* DnaN with bound CBM peptide QLDLF (PDB: 3Q4J). Subsites I and II of each binding pocket are highlighted in orange and teal, respectively. (B) Zoomed-in images of the (left) *B. subtilis* DnaN (right) and *E. coli* DnaN binding pockets. FP response for (C) PolC and (D) DnaE CBM peptides with (left) *B. subtilis* DnaN and (right) *E. coli* DnaN. Error bars show standard deviation of at least two replicates. Fits to a 1:1 binding model are represented as solid lines; fits extending past the highest protein concentration tested are represented as dashed lines.

Polymerases and other binding partners bind the clamp at a common binding site (Figure 1B), one on each DnaN protomer, through short peptide sequences called clamp-binding motifs (CBMs).^4,6^ CBMs are highly conserved across different bacterial species. An initial study identified a conserved pentapeptide CBM, with the general sequence QφxLF, with a preference for an aliphatic residue (φ) at position two, and the specific consensus sequence QL[S/D]LF.^6^ Subsequently, the hexapeptide CBM sequence QφxLxL was also identified in several *E. coli* proteins.^4,7–9^ The clamp binding pocket has two subsites; in general, the C-terminal residues (LF) bind a deep cleft in subsite I and the N-terminal glutamine Q binds in subsite II (Figure 1B).^4,9^

Most studies of clamp binding have been performed with the *E. coli* clamp. Clamp binding affinities have been measured using a range of different techniques, including fluorescence polarization (FP),^10,11^ surface plasmon resonance (SPR),^12,13^ and isothermal titration calorimetry (ITC).^10,13^ The use of CBM peptides instead of full-length proteins has been shown generally to recapitulate binding in these assays, although flanking residues may contribute to binding in some cases.^4^ In *E. coli*, typical dissociation constant (*K*_D_) values for clamp-CBM peptide binding are on the order of approximately 200 nM to 5 µM, depending on the CBM sequence.^4,10,12^ Although clamp binding pockets are highly conserved, there are still differences across species.^4,13^ Wolff et al. identified binding site differences in clamps from several bacterial species, including *E. coli* and *B. subtilis*, and reported some variation in binding affinities of several model CBMs for gram-negative vs. gram-positive clamps.^13^ However, relatively little work has been done to compare clamp binding in bacterial species beyond *E. coli*.

In a previous study, we found that the binding affinity of a consensus CBM (QLSLF) was in the mid-micromolar range for the *B. subtilis* clamp, five-fold weaker than its reported affinity for the *E. coli* clamp.^14^ This finding motivated us to explore clamp binding in *B. subtilis* more systematically. In this study, we use FP binding assays to quantify clamp binding by a range of CBM peptides using both *E. coli* and *B. subtilis* clamps. We test several reported tight-binding peptides derived from *E. coli* proteins, as well as a selection of CBMs from endogenous *B. subtilis* clamp-binding proteins. We use alanine scanning mutagenesis to show that the same CBM residues are important for binding to the *B. subtilis* and *E. coli* clamps, though the affinities are generally lower for the *B. subtilis* clamp. Notably, we find that hexapeptide CBMs bind significantly more weakly to the *B. subtilis* clamp than to the *E. coli* clamp, suggesting that there may be species-specific differences in clamp-CBM interactions.

We followed our previous approach to quantify binding of CBM peptides to the *B. subtilis* and *E. coli* clamps using FP (see Supporting Information for detailed experimental methods).^14^ In brief, we synthesized CBM peptides with the central pentapeptide or hexapeptide motif flanked on each side by two or three amino acid residues from the native protein sequence. Peptides were amidated at the C-terminus, except for CBMs that appear at the C-terminus of their respective proteins, in which case the native carboxylate group was retained. Peptides were labeled with a fluorescein moiety at the N-terminus with a 6-aminohexanoic (Ahx) acid linker. Peptide purity was determined using mass spectrometry (Table S1) and analytical HPLC (Figure S1). We performed direct binding FP assays using a fixed concentration of peptide (20 nM) and a range of clamp concentrations (three-fold dilutions starting from 50 µM for the *B. subtilis* clamp and 10 µM for the *E. coli* clamp). We fit the resulting FP response vs. clamp concentration curves to determine the dissociation constant (*K*_D_) values as described in the Supporting Information.^15^ All fit parameters are reported in Table 1 and Table S2, and figure data have been deposited in a Zenodo repository (DOI: 10.5281/zenodo.20413774).

**Table 1.** CBM peptides and measured *K*_D_ values for *B. subtilis* DnaN and *E. coli* DnaN.

| Source Protein | CBM Sequence | *B. subtilis* $K_D$ (μM) | *E. coli* $K_D$ (μM) | Ratio of *B. subtilis* to *E. coli* $K_D$ |
| --- | --- | --- | --- | --- |
| *B. subtilis* PolC | DQN-QLSLF-OH | $3.1 \pm 0.5$ | $0.64 \pm 0.01$ | 4.8 |
| *B. subtilis* DnaE | DD-QMGLF-LDE | $56 \pm 9$ | $23 \pm 2$ | 2.4 |
| *B. subtilis* Pol Y1 | YK-QLDLF-SFN | $0.9 \pm 0.1$ | $0.4144 \pm 0.0005$ | 2 |
| *B. subtilis* Pol Y2 | DDIW-QLNLF-QD | $4 \pm 1$ | $1.48 \pm 0.08$ | 2 |
| *B. subtilis* MutS | PA-QLSFF-DEA | $5.73 \pm 0.04$ | $1.79 \pm 0.08$ | 3.20 |
| *B. subtilis* MutL | EV-QEMIVP-LT | No binding | No binding | N/A |
| *B. subtilis* Pol I | ND-QLELF-EE | $13 \pm 1$ | $2.17 \pm 0.05$ | 5.8 |
| *E. coli* Pol III | SE-QVELEF-D | $26 \pm 7$ | $2.5 \pm 0.4$ | 10 |
| *E. coli* Pol IV | R-QLVLGL-OH | $8.9 \pm 0.2$ | $0.393 \pm 0.003$ | 23 |
| *E. coli* Hda | PA-QLSLPL-YL | $5 \pm 1$ | $0.44 \pm 0.02$ | 10 |

First, we characterized binding of the CBMs of the *B. subtilis* replicative polymerases PolC (QLSLF) and DnaE (QMGLF), which were the focus of our prior study.^14^ Consistent with our previous report, we found that the highly conserved PolC CBM bound with mid-micromolar affinity (Figure 1C; *K*_D_ = 3.1 ± 0.5 µM) to the *B. subtilis* clamp, whereas the slightly less conserved DnaE CBM bound with low affinity (Figure 1D; *K*_D_ > 50 µM). Next, we tested binding of these two CBMs to the *E. coli* clamp. We found that the PolC CBM bound five-fold more tightly (Figure 1C; *K*_D_ = 0.64 ± 0.01 µM), in excellent agreement with a prior study.^12^ The DnaE CBM bound more tightly than in *B. subtilis*, though still relatively weakly (Figure 1D; *K*_D_ > 10 µM). These initial results suggested that clamp binding may be generally weaker for the *B. subtilis* clamp compared to the *E. coli* clamp.

Next, we asked whether differences in binding between the two species could reflect differing importance of the five residues in the pentapeptide CBM. We used alanine scanning mutagenesis to replace each residue in the consensus motif QLDLF with an alanine and then assayed binding to the *B. subtilis* and *E. coli* clamps (Figure S2). To match a prior study, these peptides contained only the core CBM motif without flanking residues.^10^ Previous work in *E. coli* has shown that the first, fourth, and fifth CBM positions are most important for binding.^6,10,12,16^ Consistent with these prior results, we found that the core CBM peptide bound tightly (*K*_D_ = 0.103 ± 0.005 µM) and mutating the first, fourth, or fifth residues to alanine significantly weakened binding in to the *E. coli* clamp. Likewise, we found that mutating the second residue to alanine weakened binding though less substantially, and mutating the third residue had no impact.

We then tested binding of these peptides to the *B. subtilis* clamp. Because stocks of purified clamp were limited, we used a different clamp preparation for the alanine scan assays than we did in other experiments, first validating that binding was similar, albeit not quantitatively identical, for the two batches of protein (Figure S3 and Table S3). Results for the *B. subtilis* clamp were generally consistent with those for the *E. coli* clamp, with mutation of the first, fourth, or fifth residues eliminating binding (Figure S2). However, in contrast to *E. coli*, we found that the core QLDLF motif bound relatively weakly (*K*_D_ = 28 ± 8 µM), whereas the peptide with alanine substituted at the third residue bound more tightly (*K*_D_ = 7.9 ± 0.3 µM). Consistent with results for the *E. coli* clamp, mutation of the second residue to alanine weakened binding, although the effect was larger. Taken together, these data indicate that the same pentapeptide CBM positions are important for binding to the *B. subtilis* and *E. coli* clamps.

To compare clamp binding between *B. subtilis* and *E. coli* in greater detail, we chose to test five additional CBMs, four pentapeptides and one hexapeptide, derived from native *B. subtilis* proteins. Four of these CBMs were identified previously: those from the translesion synthesis (TLS) polymerases Pol Y1 (QLDLF) and Pol Y2 (QLNLF)^17,18^ and the mismatch repair (MMR) factors MutS (QLSFF)^19^ and MutL (QEMIVP).^20^ We identified one additional likely pentapeptide CBM in a protein known to bind the clamp in *E. coli*, the replication and repair polymerase Pol I (QLELF). In FP assays, we found that all four pentapeptide CBMs (Pol Y1, Pol Y2, MutS, and Pol I) bound to both the *B. subtilis* and *E. coli* clamps. For these four CBMs, binding was weaker in *B. subtilis* by a factor of two-to six-fold. As expected, the consensus Pol Y1 CBM (Figure 2A) had the highest affinity, whereas deviations from the consensus motif in the other CBMs led to weaker binding. For example, the switch from the consensus, negatively-charged aspartic acid in the third position in the Pol Y1 CBM (QLDLF) to the sterically similar but neutral asparagine in the Pol Y2 CBM (QLNLF) (Figure 2B) weakened binding by 3 – 4-fold in both species, suggesting a role for the negative charge in binding affinity. Prior studies with the *E. coli* clamp have reported similar fold-changes in affinity for this or similar substitutions.^10,12^ Switching the aspartic acid to glutamic acid in the Pol I CBM (QLELF) (Figure 2E), which conserves the negative charge but changes the residue’s shape and size, also led to a similar decrease in affinity. Differences in other conserved residues also affected binding affinity. In particular, switching the highly conserved leucine at the fourth position in the PolC CBM (QLSLF) to phenylalanine instead in the MutS CBM (QLSFF) (Figure 2C) produced a 1.5 – 3-fold decrease in affinity.

**Figure 2.**
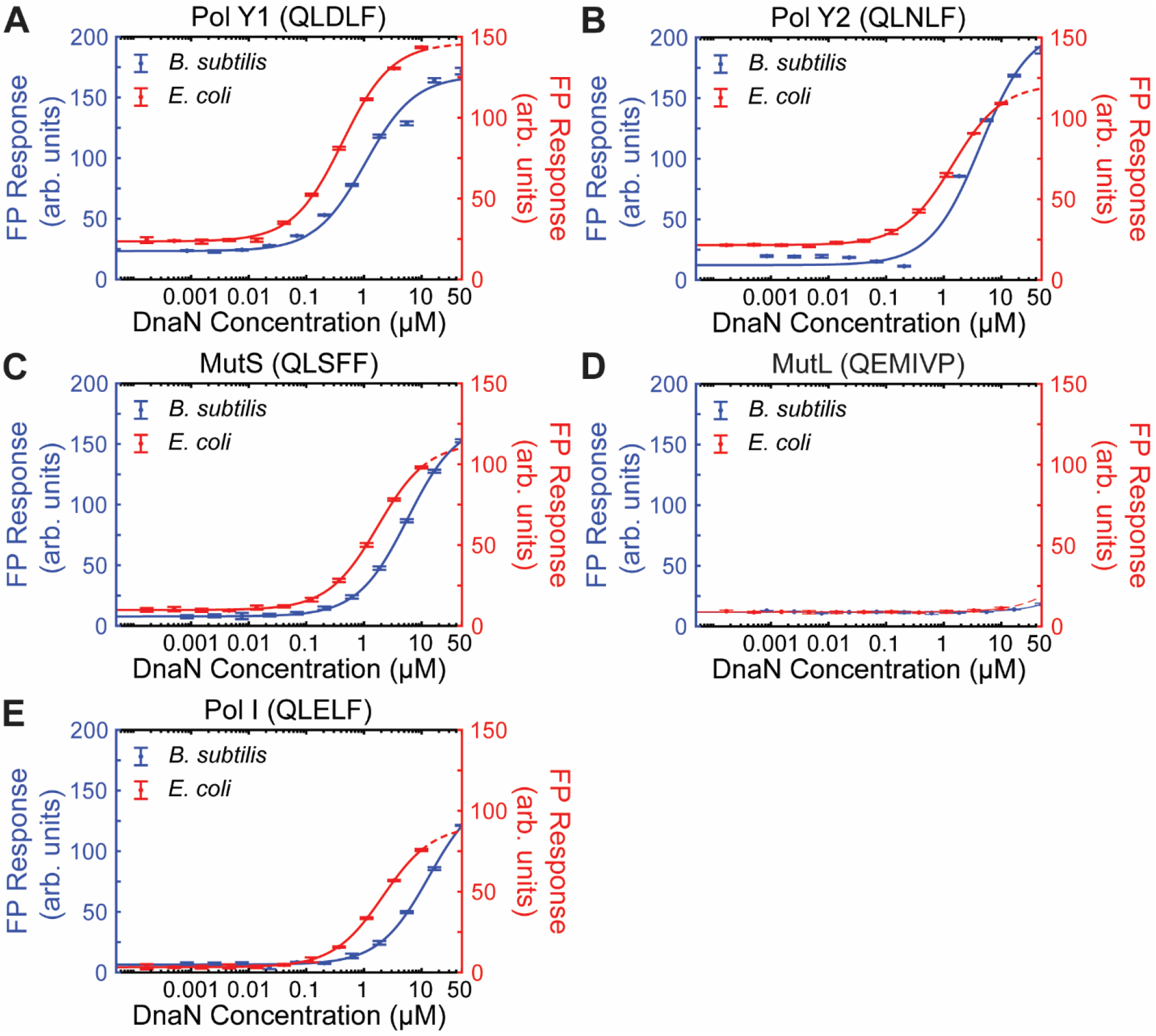
FP response for (A) Pol Y1, (B) Pol Y2, (C) MutS, (D) MutL, and (E) Pol I CBM peptides with (left) *B. subtilis* DnaN and (right) *E. coli* DnaN. Error bars show standard deviation of at least two replicates. Fits to a 1:1 binding model are represented as solid lines; fits extending past the highest protein concentration tested are represented as dashed lines.

Surprisingly, the MutL (Figure 2D) CBM did not bind measurably to either clamp. The MutL CBM (QEMIVP) does not agree well with the consensus pentapeptide or hexapeptide motifs, containing a charged glutamic acid in the second position, instead of an aliphatic amino acid, and lacking the highly conserved leucine and/or phenylalanine residues in the fourth through sixth positions. Binding was observed previously in a crystal structure in which the MutL regulatory domain was fused to the full-length clamp protein; it is possible that tethering enhanced what is naturally a weak interaction by increasing the local concentration of the MutL CBM near the clamp binding pocket.^20^

In addition to the conserved pentapeptide CBM motif, a hexapeptide (QφxLxL) motif has also been identified in several *E. coli* proteins.^4^ However, we did not observe binding for the MutL CBM, which is (to our knowledge) the only hexapeptide CBM identified in *B. subtilis* proteins. Thus, we asked whether other hexapeptide CBMs from *E. coli* would bind to the *B. subtilis* clamp. We tested three hexapeptide CBMs derived from native *E. coli* proteins: the C-terminal CBM from the α subunit of the replicative polymerase Pol III (QVELEF), the CBM from the TLS polymerase Pol IV (QLVLGL), and the CBM from the replication initiation regulator Hda (QLSLPL). All three sequences have been identified as tight-binding CBMs in *E. coli*, with reported *K*_D_ values in the sub-micromolar range. In agreement with these prior reports, we found that the Pol IV (Figure 3B; *K*_D_ = 0.393 ± 0.003 µM)^9^ and Hda (Figure 3C; *K*_D_ = 0.44 ± 0.02 µM)^8^ CBMs bound tightly in *E. coli*. Binding was somewhat weaker for the Pol III CBM (Figure 3A; *K*_D_ = 2.5 ± 0.4 µM), also consistent with previous studies.^7^ Interestingly, although all three CBM peptides also bound to the *B. subtilis* clamp, binding was roughly 10-to 20-fold weaker: Pol IV (Figure 3B; *K*_D_ = 8.9 ± 0.02 µM), Hda (Figure 3C; *K*_D_ = 5 ± 1 µM), Pol III (Figure 3A; *K*_D_ = 26 ± 7 µM). We note that our results for the Pol IV CBM are in disagreement with a prior study that reported similar binding affinities for this peptide in both *E. coli* and *B. subtilis*, measured via SPR.^13^ Our results show that several known hexapeptide CBMs bind more weakly in *B. subtilis* than in *E. coli*, suggesting that the utilization of different CBM lengths may be species-specific.

**Figure 3.**
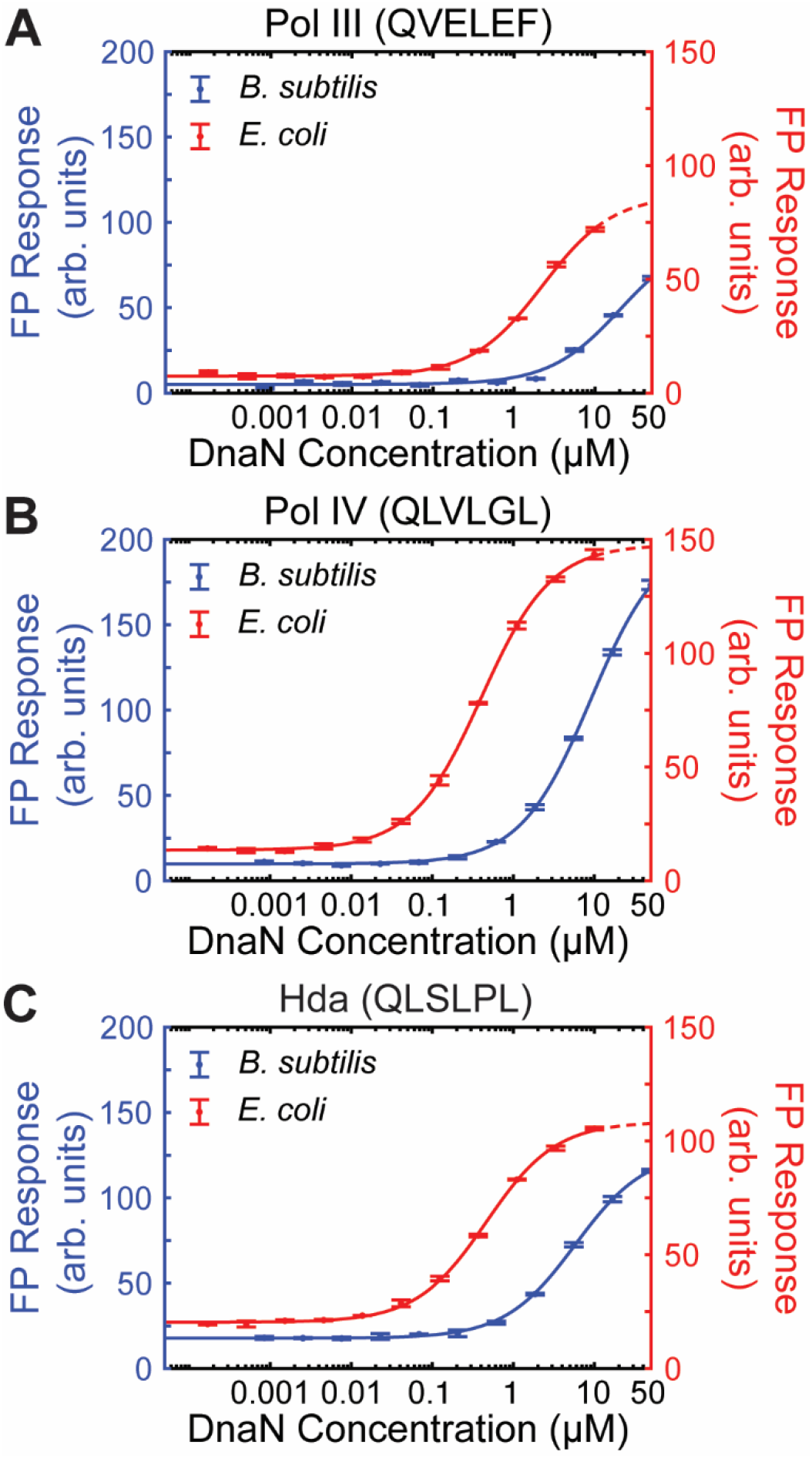
FP response for (A) Pol III, (B) Pol IV, and (C) Hda CBM peptides with *B. subtilis* DnaN and *E. coli* DnaN. Error bars show standard deviation of at least two replicates. Fits to a 1:1 binding model are represented as solid lines; fits extending past the highest protein concentration tested are represented as dashed lines.

In conclusion, we have systematically compared clamp binding in the model gram-negative bacterium *E. coli* and the model gram-positive bacterium *B. subtilis* for a range of pentapeptide and hexapeptide CBMs using FP binding assays. As expected, we find similarities between the two species, including agreement in the importance of the different amino acid positions in the conserved pentapeptide motif QLDLF. However, our results also reveal differences in clamp binding between the two species. Most notably, we find that, although pentapeptide CBMs bind somewhat more weakly to the *B. subtilis* clamp compared to the *E. coli* clamp, hexapeptide CBMs bind at least an order of magnitude more weakly. We also find that the only reported endogenous hexapeptide CBM in *B. subtilis* does not bind over the concentration range tested. Taken together, our results suggest that hexapeptide CBMs may be less universal than pentapeptide CBMs, although systematic bioinformatics analysis would be required to confirm or deny this hypothesis. Future work should explore the biological causes and consequences of differences in hexapeptide CBM affinity across bacterial species, as such differences may also represent an opportunity for the design of species-selective drugs or probes that target the clamp.

## ASSOCIATED CONTENT

### Supporting Information

The following files are available free of charge.

Detailed experimental procedures, peptide characterization data, and additional FP binding assay data (PDF)

## AUTHOR INFORMATION

### Authors

**Sarah J. Rancic –** *Department of Chemistry and Biochemistry, Fordham University, Bronx, NY 10458, United States*;

**Katrin M. Klassen –** *Department of Chemistry and Biochemistry, Fordham University, Bronx, NY 10458, United States*;

### Author Contributions

The manuscript was written through contributions of all authors. All authors have given approval to the final version of the manuscript.

## ACKNOWLEDGMENTS

We thank Luke O’Neal for assistance with figure preparation. This work was supported by the National Institute of General Medical Sciences of the National Institutes of Health [award number R15GM151677 to E.S.T.]. Additional support was provided by the Fordham College at Rose Hill Undergraduate Research Grant program [award to K.M.K.]. Data were acquired on a BioTek Synergy H1MF plate reader, generously donated to the Department of Chemistry and Biochemistry by Dr. Michael B. Dowell.

## ABBREVIATIONS

CBM: clamp-binding motif
FP: fluorescence polarization.

## REFERENCES

(1) Johnson, A.; O’Donnell, M. Cellular DNA Replicases: Components and Dynamics at the Replication Fork. Annu Rev Biochem 2005, 74, 283–315. 10.1146/annurev.biochem.73.011303.073859.

(2) McHenry, C. S. DNA Replicases from a Bacterial Perspective. Annu. Rev. Biochem. 2011, 80 (1), 403–436. 10.1146/annurev-biochem-061208-091655.

(3) Altieri, A. S.; Kelman, Z. DNA Sliding Clamps as Therapeutic Targets. Front. Mol. Biosci. 2018, 5, 87. 10.3389/fmolb.2018.00087.

(4) Simonsen, S.; Søgaard, C. K.; Olsen, J. G.; Otterlei, M.; Kragelund, B. B. The Bacterial DNA Sliding Clamp, β-Clamp: Structure, Interactions, Dynamics and Drug Discovery. Cell. Mol. Life Sci. 2024, 81 (1), 245. 10.1007/s00018-024-05252-w.

(5) Mulye, M.; Singh, M. I.; Jain, V. From Processivity to Genome Maintenance: The Many Roles of Sliding Clamps. Genes 2022, 13 (11), 2058. 10.3390/genes13112058.

(6) Dalrymple, B. P.; Kongsuwan, K.; Wijffels, G.; Dixon, N. E.; Jennings, P. A. A Universal Protein–Protein Interaction Motif in the Eubacterial DNA Replication and Repair Systems. Proc. Natl. Acad. Sci. U.S.A. 2001, 98 (20), 11627–11632. 10.1073/pnas.191384398.

(7) López De Saro, F. J.; Georgescu, R. E.; O’Donnell, M. A Peptide Switch Regulates DNA Polymerase Processivity. Proc. Natl. Acad. Sci. U.S.A. 2003, 100 (25), 14689–14694. 10.1073/pnas.2435454100.

(8) Kurz, M.; Dalrymple, B.; Wijffels, G.; Kongsuwan, K. Interaction of the Sliding Clamp Beta-Subunit and Hda, a DnaA-Related Protein. J Bacteriol 2004, 186 (11), 3508–3515. 10.1128/JB.186.11.3508-3515.2004.

(9) Burnouf, D. Y.; Olieric, V.; Wagner, J.; Fujii, S.; Reinbolt, J.; Fuchs, R. P. P.; Dumas, P. Structural and Biochemical Analysis of Sliding Clamp/Ligand Interactions Suggest a Competition Between Replicative and Translesion DNA Polymerases. Journal of Molecular Biology 2004, 335 (5), 1187–1197. 10.1016/j.jmb.2003.11.049.

(10) Yin, Z.; Kelso, M. J.; Beck, J. L.; Oakley, A. J. Structural and Thermodynamic Dissection of Linear Motif Recognition by the E. Coli Sliding Clamp. J. Med. Chem. 2013, 56 (21), 8665– 8673. 10.1021/jm401118f.

(11) Georgescu, R. E.; Yurieva, O.; Kim, S.-S.; Kuriyan, J.; Kong, X.-P.; O’Donnell, M. Structure of a Small-Molecule Inhibitor of a DNA Polymerase Sliding Clamp. Proc. Natl. Acad. Sci. U.S.A. 2008, 105 (32), 11116–11121. 10.1073/pnas.0804754105.

(12) Wijffels, G.; Dalrymple, B. P.; Prosselkov, P.; Kongsuwan, K.; Epa, V. C.; Lilley, P. E.; Jergic, S.; Buchardt, J.; Brown, S. E.; Alewood, P. F.; Jennings, P. A.; Dixon, N. E. Inhibition of Protein Interactions with the β_2_ Clamp of Escherichia Coli DNA Polymerase III by Peptides from β_2_-Binding Proteins. Biochemistry 2004, 43 (19), 5661–5671. 10.1021/bi036229j.

(13) Wolff, P.; Amal, I.; Oliéric, V.; Chaloin, O.; Gygli, G.; Ennifar, E.; Lorber, B.; Guichard, G.; Wagner, J.; Dejaegere, A.; Burnouf, D. Y. Differential Modes of Peptide Binding onto Replicative Sliding Clamps from Various Bacterial Origins. J. Med. Chem. 2014, 57 (18), 7565–7576. 10.1021/jm500467a.

(14) O’Neal, L. G.; Drucker, M. N.; Lai, N. K.; Clemente, A. F.; Campbell, A. P.; Way, L. E.; Hong, S.; Holmes, E. E.; Rancic, S. J.; Sawyer, N.; Wang, X.; Thrall, E. S. The B. Subtilis Replicative Polymerases Bind the Sliding Clamp with Different Strengths to Tune Their Activity in DNA Replication. Nucleic Acids Research 2025, 53 (14), gkaf721. 10.1093/nar/gkaf721.

(15) Jarmoskaite, I.; AlSadhan, I.; Vaidyanathan, P. P.; Herschlag, D. How to Measure and Evaluate Binding Affinities. eLife 2020, 9, e57264. 10.7554/eLife.57264.

(16) Wijffels, G.; Johnson, W. M.; Oakley, A. J.; Turner, K.; Epa, V. C.; Briscoe, S. J.; Polley, M.; Liepa, A. J.; Hofmann, A.; Buchardt, J.; Christensen, C.; Prosselkov, P.; Dalrymple, B. P.; Alewood, P. F.; Jennings, P. A.; Dixon, N. E.; Winkler, D. A. Binding Inhibitors of the Bacterial Sliding Clamp by Design. J. Med. Chem. 2011, 54 (13), 4831–4838. 10.1021/jm2004333.

(17) Duigou, S.; Ehrlich, S. D.; Noirot, P.; Noirot-Gros, M.-F. Distinctive Genetic Features Exhibited by the Y-Family DNA Polymerases in Bacillus Subtilis. Mol Microbiol 2004, 54 (2), 439–451. 10.1111/j.1365-2958.2004.04259.x.

(18) Marrin, M. E.; Foster, M. R.; Santana, C. M.; Choi, Y.; Jassal, A. S.; Rancic, S. J.; Greenwald, C. R.; Drucker, M. N.; Feldman, D. T.; Thrall, E. S. The Translesion Polymerase Pol Y1 Is a Constitutive Component of the B. Subtilis Replication Machinery. Nucleic Acids Research 2024, 52 (16), 9613–9629. 10.1093/nar/gkae637.

(19) Simmons, L. A.; Davies, B. W.; Grossman, A. D.; Walker, G. C. β Clamp Directs Localization of Mismatch Repair in Bacillus Subtilis. Molecular Cell 2008, 29 (3), 291–301. 10.1016/j.molcel.2007.10.036.

(20) Almawi, A. W.; Scotland, M. K.; Randall, J. R.; Liu, L.; Martin, H. K.; Sacre, L.; Shen, Y.; Pillon, M. C.; Simmons, L. A.; Sutton, M. D.; Guarné, A. Binding of the Regulatory Domain of MutL to the Sliding β-Clamp Is Species Specific. Nucleic Acids Research 2019, 47 (9), 4831–4842. 10.1093/nar/gkz115.

