## Supplementary Information for "Characterizing the Molecular Determinants of Clamp Binding in *B. subtilis*"

**This PDF file includes:**

Figures S1 to S3
Tables S1 to S3
Supplementary Methods
Supplementary References

### Supplementary Figures

*B. subtilis* Pol Y1 (SR2A)

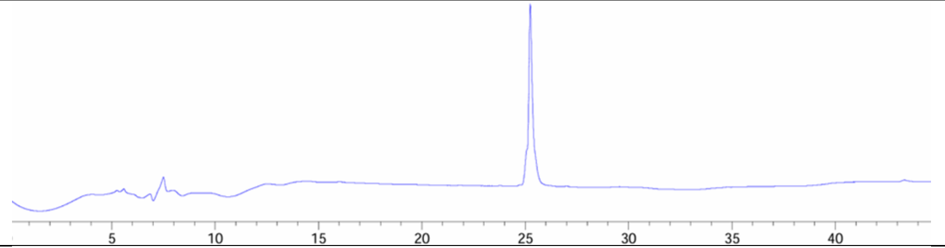


*B. subtilis* Pol Y2 (SR8A)

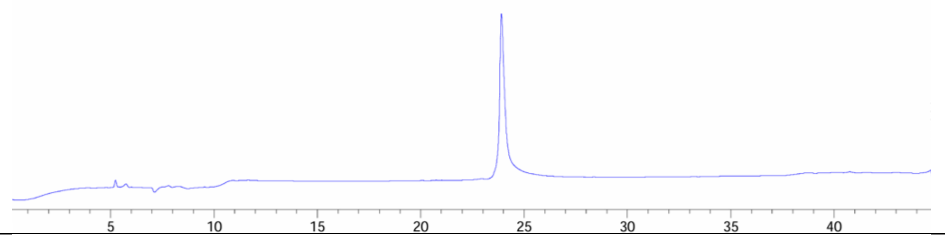


*B. subtilis* MutS (SR4A)

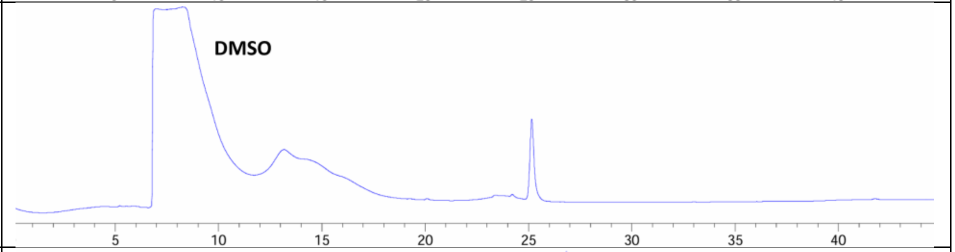


*B. subtilis* MutL (SR5A)

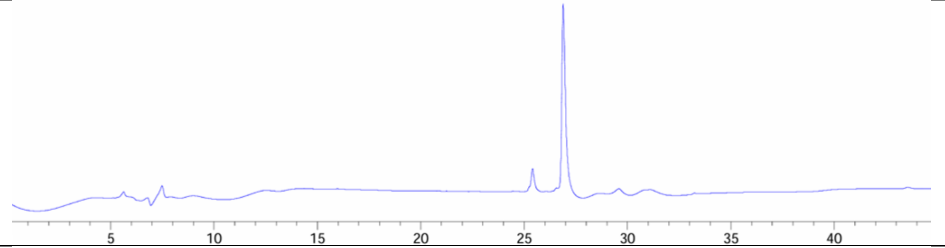


*B. subtilis* Pol I (SR10A)

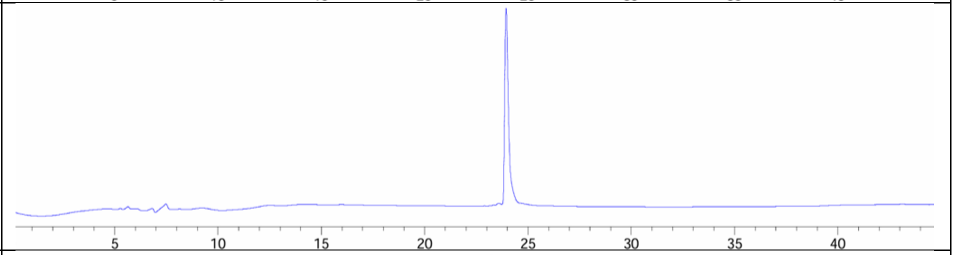


*E. coli* Pol III

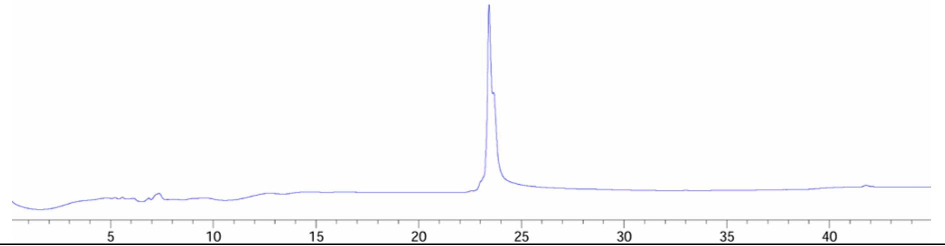


**Figure S1.** Analytical HPLC chromatograms for all new peptides, measured at 220 nm.

| | |
|---|---|
| *E. coli* Pol IV (SR1A) | 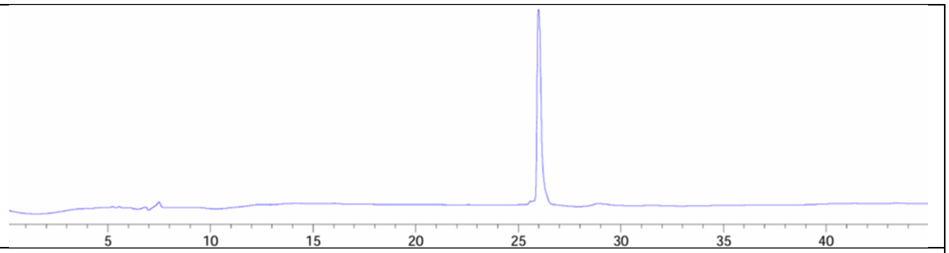  |
| QLDLF (SR12A) | 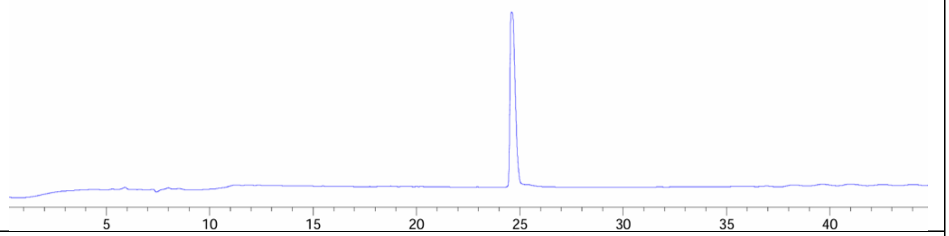  |
| ALDLF (SR13A) | 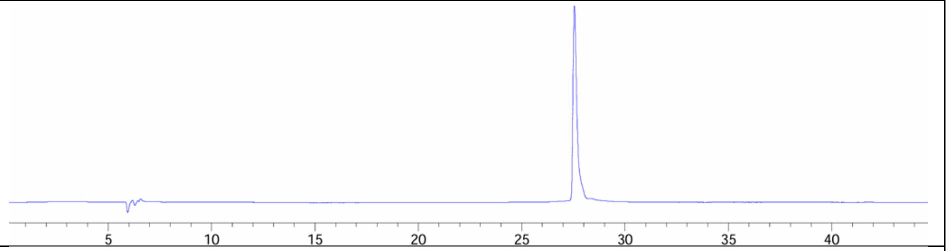  |
| QADLF (SR14A) | 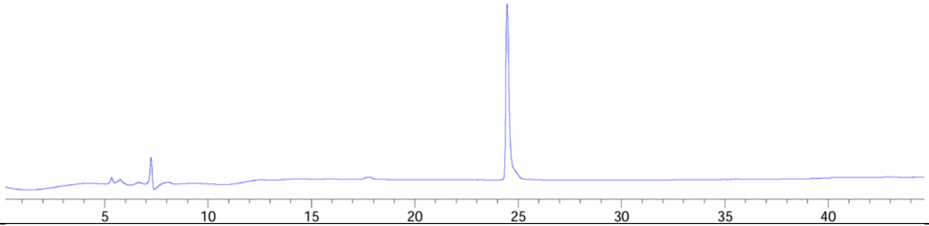  |
| QLALF (SR15A) | 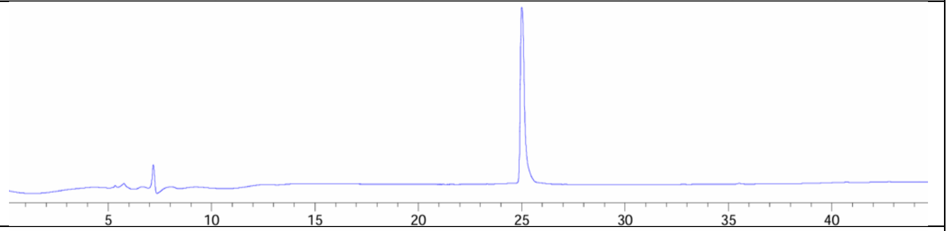  |
| QLDAF (SR16A) | 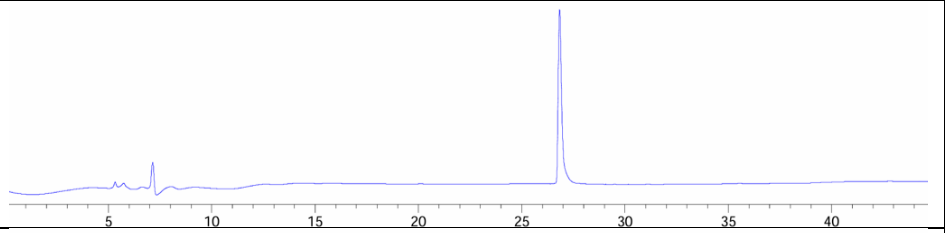  |
| QLDLA (SR17A) | 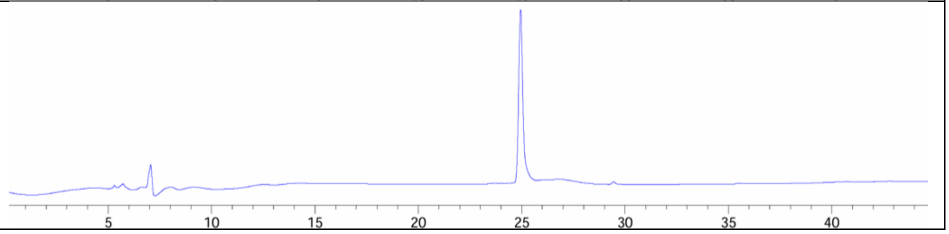  |

**Figure S1.** Analytical HPLC chromatograms for all new peptides, measured at 220 nm (cont.).

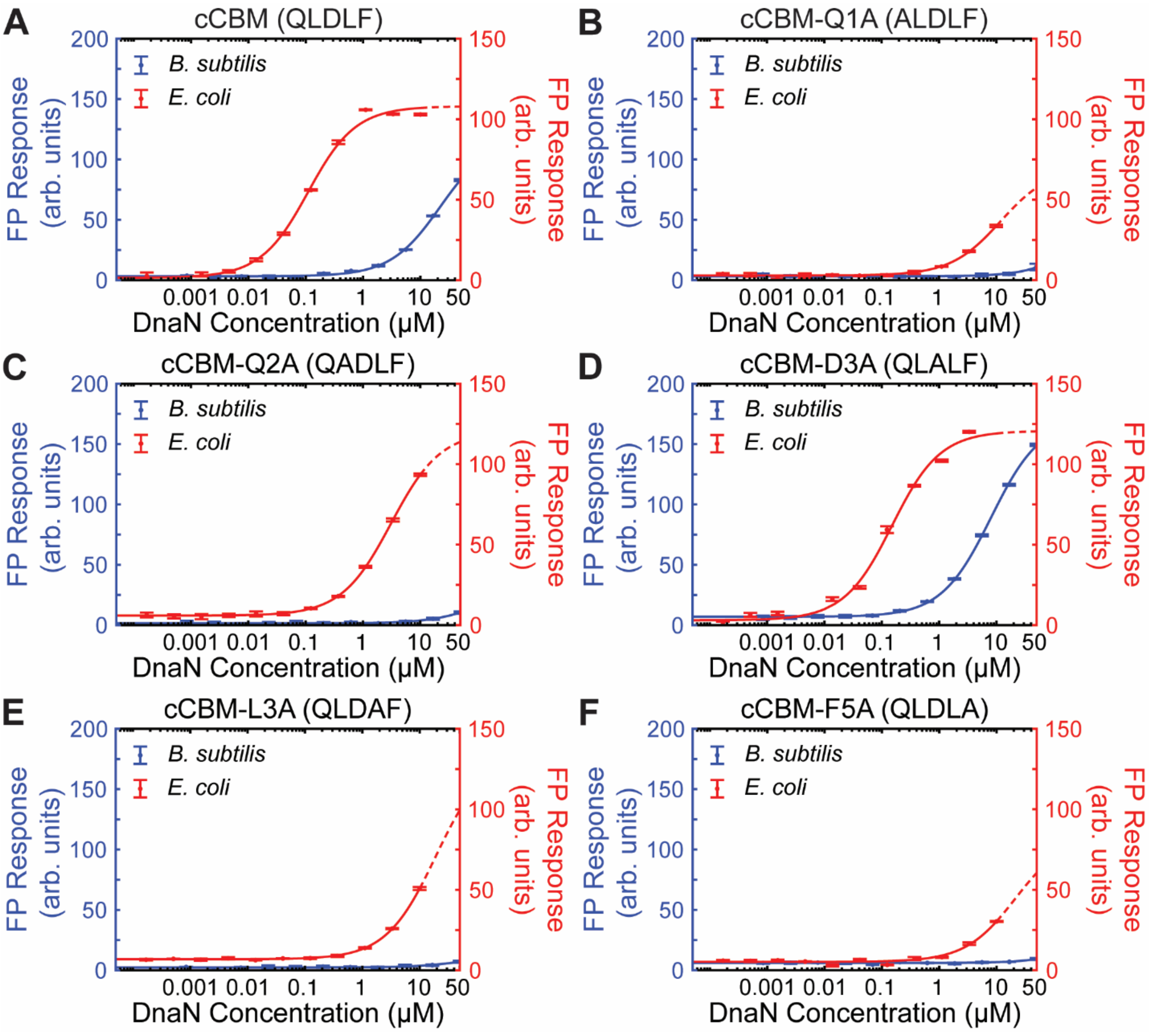


**Figure S2.** CBM alanine scan binding data. FP response for (A) cCBM (QLDLF), (B) cCBM-Q1A (ALDLF), (C) cCBM-L2A (QADLF), (D) cCBM-D3A (QLALF), (E) cCBM-L4A (QLDAF), and (F) cCBM-F5A (QLDLA) CBM peptides with (left) *B. subtilis* DnaN and (right) *E. coli* DnaN. Error bars show standard deviation of at least two replicates. Fits to a 1:1 binding model are represented as solid lines; fits extending past the highest protein concentration tested are represented as dashed lines.

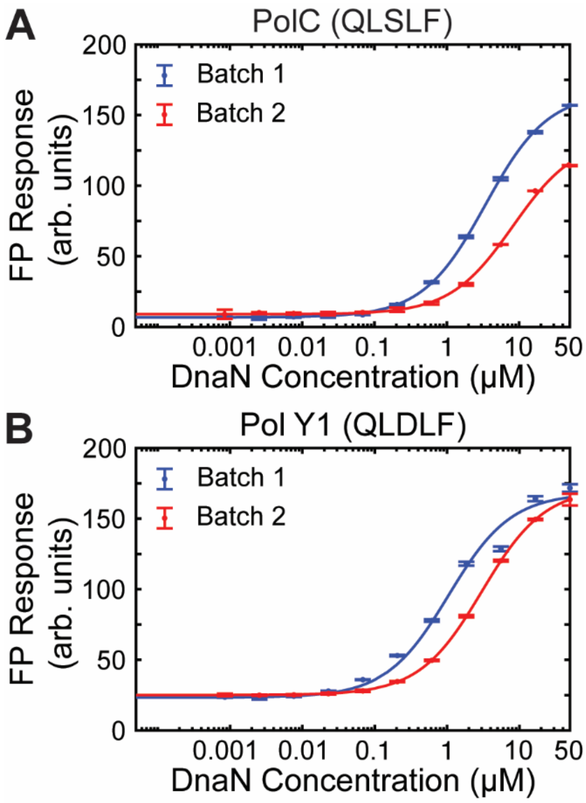


**Figure S3.** Cross-validation of *B. subtilis* DnaN protein batches. FP response for (A) PolC (QLSLF) and (B) Pol Y1 (QLDLF) CBM peptides with primary DnaN batch #1 (data in Tables 1 and S3) and secondary DnaN batch #2 used for alanine scan assays only (data in Table S3). Error bars show standard deviation of at least two replicates. Fits to a 1:1 binding model are represented as solid lines.

### Supplementary Tables

**Table S1.** MALDI mass spectrometry characterization of new CBM peptides. FITC = fluorescein-5-thiocarbamoyl, FAM = fluorescein-5(6)-carbonyl, Ahx = 6-aminohexanoyl; -OH indicates a C-terminal carboxylic acid; -$NH_2$ indicates a C-terminal amide. Clamp binding motif (CBM) sequences are underlined.

| Source Protein | Peptide Name | Full Sequence | Expected *m/z* $(M+H^+)$ | Observed *m/z* $(M+H^+)$ |
|---|---|---|---|---|
| *B. subtilis* Pol Y1 | SR2A | FAM-Ahx-YKQLDLFSFN-$NH_2$ | 1744.8 | 1744.7 |
| *B. subtilis* Pol Y2 | SR8A | FAM-Ahx-DDIWQLNLFQD-$NH_2$ | 1877.8 | 1877.6 |
| *B. subtilis* MutS | SR4A | FAM-Ahx-PAQLSFFDEA-$NH_2$ | 1594.7 | 1594.3 |
| *B. subtilis* MutL | SR5A | FAM-Ahx-EVQEMIVPLT-$NH_2$ | 1628.8 | 1629.9 |
| *B. subtilis* Pol I | SR10A | FAM-Ahx-NDQLELFEE-$NH_2$ | 1606.7 | 1606.7 |
| *E. coli* Pol III | Pol III | FAM-Ahx-SEQVELEFD-OH | 1566.6 | 1566.4 |
| *E. coli* Pol IV | SR1A | FAM-Ahx-RQLVLGL-OH | 1269.6 | 1269.5 |
| Consensus CBM (cCBM) | SR12A | FAM-Ahx-QLDLF-OH | 1106.4 | 1107.4 |
| cCBM-Q1A | SR13A | FAM-Ahx-ALDLF-OH | 1049.4 | 1049.4 |
| cCBM-L2A | SR14A | FAM-Ahx-QADLF-OH | 1064.4 | 1064.5 |
| cCBM-D3A | SR15A | FAM-Ahx-QLALF-OH | 1062.5 | 1062.8 |
| cCBM-L4A | SR16A | FAM-Ahx-QLDAF-OH | 1064.4 | 1065.1 |
| cCBM-F5A | SR17A | FAM-Ahx-QLDLA-OH | 1030.4 | 1030.4 |

**Table S2.** Complete fit parameters for CBM peptides with *B. subtilis* DnaN and *E. coli* DnaN. $K_D$: dissociation constant in µM. *A*: amplitude (arb. units). *B*: baseline (arb. units). Values represent mean ± standard deviation.

| **Source Protein** | **CBM Sequence** | **$K_D$ (µM)** | | ***A*** | | ***B*** | |
|---|---|---|---|---|---|---|---|
| | | ***B. subtilis*** | ***E. coli*** | ***B. subtilis*** | ***E. coli*** | ***B. subtilis*** | ***E. coli*** |
| *B. subtilis* PolC | DQN-<u>QLSLF</u>-OH | 3.1 ± 0.5 | 0.64 ± 0.01 | 150 ± 10 | 104 ± 2 | 7.1 ± 0.5 | 7.5 ± 0.6 |
| *B. subtilis* DnaE | DD-<u>QMGLF</u>-LDE | 56 ± 9 | 23 ± 2 | 140 ± 30 | 76 ± 4 | 11.6 ± 0.8 | 12 ± 1 |
| *B. subtilis* Pol Y1 | YK-<u>QLDLF</u>-SFN | 0.9 ± 0.1 | 0.4144 ± 0.0005 | 143 ± 3 | 120 ± 4 | 24 ± 1 | 24 ± 1 |
| *B. subtilis* Pol Y2 | DDIW-<u>QLNLF</u>-QD | 4 ± 1 | 1.48 ± 0.08 | 190 ± 10 | 101 ± 2 | 14 ± 2 | 21 ± 1 |
| *B. subtilis* MutS | PA-<u>QLSFF</u>-DEA | 5.73 ± 0.04 | 1.79 ± 0.08 | 161.011 ± 0.005 | 103 ± 1 | 9 ± 1 | 9.79 ± 0.09 |
| *B. subtilis* MutL | EV-<u>QEMIVP</u>-LT | No binding | No binding | N/A | N/A | N/A | N/A |
| *B. subtilis* Pol I | ND-<u>QLELF</u>-EE | 13 ± 1 | 2.17 ± 0.05 | 143 ± 3 | 87 ± 1 | 6.6 ± 0.1 | 5 ± 2 |
| *E. coli* Pol III | SE-<u>QVELEF</u>-D | 26 ± 7 | 2.5 ± 0.4 | 100 ± 20 | 86 ± 9 | 6 ± 2 | 6 ± 2 |
| *E. coli* Pol IV | R-<u>QLVLGL</u>-OH | 8.9 ± 0.2 | 0.393 ± 0.003 | 188 ± 6 | 130 ± 10 | 10.4 ± 0.6 | 13.1 ± 0.7 |
| *E. coli* Hda | PA-<u>QLSLPL</u>-YL | 5 ± 1 | 0.44 ± 0.02 | 90 ± 20 | 80 ± 10 | 16.9 ± 0.7 | 18 ± 3 |
| Consensus CBM (cCBM) | <u>QLDLF</u> | 28 ± 8 | 0.103 ± 0.005 | 130 ± 20 | 108 ± 2 | 5 ± 3 | 1.3 ± 0.2 |
| cCBM-Q1A | <u>ALDLF</u> | No binding | Weak binding | N/A | N/A | N/A | N/A |
| cCBM-L2A | <u>QADLF</u> | No binding | 3.4 ± 0.5 | N/A | 119 ± 6 | N/A | 7 ± 1 |
| cCBM-D3A | <u>QLALF</u> | 7.8 ± 0.3 | 0.13 ± 0.01 | 158 ± 9 | 115 ± 4 | 6.94 ± 0.04 | 5 ± 3 |
| cCBM-L4A | <u>QLDAF</u> | No binding | Weak binding | N/A | N/A | N/A | N/A |
| cCBM-F5A | <u>QLDLA</u> | No binding | Weak binding | N/A | N/A | N/A | N/A |

**Table S3.** Measured $K_D$ values for two batches of *B. subtilis* DnaN with two CBM peptides.

| **Source Protein** | **CBM Sequence** | ***B. subtilis* DnaN Batch #1 $K_D$ (µM)** | ***B. subtilis* DnaN Batch #2 $K_D$ (µM)** | **Ratio of Batch #1 to #2 $K_D$** |
|---|---|---|---|---|
| PolC | DQN-<u>QLSLF</u>-OH | 3.1 ± 0.5 | 8.4 ± 0.3 | 2.7 |
| Pol Y1 | DD-<u>QMGLF</u>-LDE | 0.9 ± 0.1 | 3.3 ± 0.3 | 4 |

### Supplementary Methods

*Peptide Synthesis and Characterization:* Peptide synthesis was performed as described previously.[1] In brief, commercial solvents and reagents were used without further purification. Nα-Fmoc-protected amino acids and other peptide synthesis reagents were purchased from Advanced ChemTech, Ambeed, ChemImpex International, Gyros Protein Technologies, and Oakwood Chemical. Peptides were synthesized following standard Fmoc solid-phase approaches starting from Rink MBHA resin (0.62 mmol/g resin) for C-terminally amidated peptides or appropriate Wang resins pre-loaded with Fmoc-Leu, Fmoc-Phe, Fmoc-Asp(OtBu), or Fmoc-Ala for peptides with C-terminal carboxylic acids (0.3 – 0.8 mmol/g resin).

Automated synthesis on a Gyros Protein Technologies PurePep Chorus synthesizer was used to introduce standard Fmoc-protected amino acids. Dried resin was initially swollen in N,N-dimethylformamide (DMF) for 10 min. Fmoc deprotection was performed by two sequential reactions of 15 min each using 20% piperidine in DMF at room temperature. For each amino acid coupling, four molar equivalents each of the Fmoc-protected amino acid and O-(1H-6-chlorobenzotriazole-1-yl)-1,1,3,3-tetramethyluronium hexafluorophosphate (HCTU) and eight molar equivalents of N-methylmorpholine (NMM) in N,N-dimethylformamide (DMF) were sequentially added to deprotected resin and reacted for 1 h at 55 °C. In between coupling and deprotection steps, the resin was washed with DMF three times.

Due to poor solubility of Fmoc-6-aminohexanoic acid (Fmoc-Ahx) in DMF, Fmoc-Ahx and 5(6)-carboxyfluorescein (FAM) were introduced manually. All Fmoc deprotection steps were performed as described above. For Fmoc-Ahx coupling, pre-activation was achieved by dissolving five molar equivalents each of Fmoc-Ahx, ethyl cyanohydroxyiminoacetate (Oxyma), and diisopropylcarbodiimide (DIC) in DMF, followed by reaction for 20 – 30 min at room temperature. The pre-activated ester was added to deprotected resin for 2 h at room temperature. To maximize coupling, this reaction was performed twice without Fmoc deprotection in between (double coupling). After the second coupling reaction, the resin was reacted with 8 molar equivalents of acetic anhydride and 1 molar equivalent of N,N-diisopropylethylamine in 2 mL DMF to add an acetyl capping group to any deprotected resin remaining. After Fmoc deprotection, FAM was pre-activated and coupled once as described for Fmoc-Ahx. In between coupling and deprotection steps, the resin was washed with DMF three times.

Periodically during synthesis, microcleavage reactions were performed by adding a small amount of resin to 1 mL of a mixture containing 2.5% $H_2O$ and 2.5% triisopropylsilane (TIPS) in trifluoroacetic acid (TFA). After 30 min, reactions were diluted with 3 mL acetonitrile and mixed 1:4 with a saturated matrix solution (α-cyano-4-hydroxycinnamic acid (CHCA) dissolved in a 1:1 mixture of acetonitrile and water + 0.1% TFA). A 1 µL volume of sample mixture was dried on a stainless steel plate and analyzed on a Shimadzu MALDI-8020 Benchtop MALDI-TOF mass spectrometer.

Peptides were globally deprotected and cleaved from dried resin by reaction with freshly prepared cleavage cocktail (92.5% TFA, 2.5% TIPS, 2.5% $H_2O$, and 2.5% 1,2-ethanedithiol (EDT)) for 2 h at room temperature. Resin beads were filtered out, the filtrate volume was reduced by rotary evaporation, and the sample was precipitated three times with ice-cold diethyl ether to yield the crude peptide.

Peptides were purified on a Shimadzu Nexera high-performance liquid chromatography (HPLC) system using gradients of water and acetonitrile (ACN) containing 0.1% trifluoroacetic acid (TFA) with preparative C18 columns. Peptide purity was evaluated on an Agilent 1100 HPLC system using a Luna 5 μm C18(2) column (150 × 4.6 mm) with a flow rate of 0.3 mL/min (gradient: 5 – 95% solvent B over 30 min, solvent A = 0.1% aqueous TFA, B = 95% ACN, 5% water, 0.1% TFA; Figure S1). High-resolution mass spectrometry data were collected on a Shimadzu MALDI-8020 Benchtop MALDI-TOF mass spectrometer using α-cyano-4-hydroxycinnamic acid as the matrix

*Protein Expression and Purification:* The purification of the N-terminally hexahistidine ($His_6$)-tagged *E. coli* DnaN (UniProt: P0A988) and *B. subtilis* DnaN (UniProt: A0A6M4JEE3) proteins used in this study was described previously.[1] In brief, pET-28b expression plasmids were transformed into *E. coli* BL21(DE3) and protein expression was induced by addition of 1 mM isopropyl β- d-1-thiogalactopyranoside (IPTG) for 3 h at 37 °C. Cell pellets were harvested by centrifugation, then resdissolved in lysis buffer (20 mM Tris-HCl (pH 7.5), 350 mM NaCl, 1 mM phenylmethylsulfonyl fluoride (PMSF), 10 mM imidazole, 5 mM β-mercaptoethanol (BME), and 1 mg/mL lysozyme) at 5 mL/g concentration and incubated at 4 °C for 30 min, followed by sonication (Fisherbrand Model 120). The lysate was cleared by centrifugation and supplemented with imidazole to a final concentration of 30 mM, then incubated with Ni-NTA agarose (Thermo Scientific HisPur resin) for 1 h at 4 °C. The column was washed sequentially with a high-salt buffer (20 mM Tris-HCl (pH 7.5), 500 mM NaCl, 5 mM BME, 1 mM PMSF, and 30 mM imidazole) and a low-salt buffer (20 mM Tris-HCl (pH 7.5), 100 mM NaCl, 5 mM BME, 1 mM PMSF, and 30 mM imidazole), after which bound protein was eluted in stages with elution buffer (20 mM Tris-HCl (pH 7.5), 100 mM NaCl, 5 mM BME, and 150 mM imidazole). Eluted protein was dialyzed overnight against dialysis buffer (20 mM Tris-HCl (pH 7.5), 100 mM NaCl, 5 mM BME, and 10% glycerol), then concentrated using a spin concentrator (Amicon Ultra-4 10,000 MWCO). Protein purity was assessed by SDS-PAGE, and the concentration was quantified spectrophotometrically using a microvolume spectrophotometer.

This study used two different preparations of *B. subtilis* DnaN. The batch used in the alanine scan FP binding assays (designated batch #2 in this study) was purified in house as described above. In addition, as described previously,[1] we purchased a custom batch of N-terminally $His_6$-tagged *B. subtilis* DnaN from a commercial source (GenScript) with the same $His_6$

tag and linker (designated batch #1 in this study). This batch was used in all other FP binding assays. Cross-validation of the two *B. subtilis* DnaN batches was performed by assaying binding to the PolC and Pol Y1 CBM peptides (Figure S2). The resulting $K_D$ values were similar, albeit with an approximately 3-fold higher value for batch #2. Thus, we did not compare quantitative results across the two batches of protein.

*Fluorescence Polarization (FP) Binding Measurements:* Binding affinities were measured using fluorescence polarization (FP) binding assays with fluorescein-tagged peptides and purified DnaN proteins exactly as described previously.[1] In brief, peptide stocks were dissolved in 10 mM sodium phosphate buffer (pH 7.4) and protein stocks were diluted in dialysis buffer; both buffers were supplemented with 0.1% Pluronic acid F-68. Proteins were dissolved to the maximum concentration used in the assay (10 µM for *E. coli* DnaN, 50 µM for *B. subtilis* DnaN) and then serially diluted 3-fold to the lowest concentration. For each protein concentration, the peptide stock was diluted 1:100 to give a final concentration of 20 nM. Samples were measured in triplicate in a black 96-well plate.

FP assays were performed on a BioTek Synergy H1MF plate reader equipped with polarization filters (excitation: 485 nm/20 nm, emission: 528 nm/20 nm). The resulting FP response vs. protein concentration curves were fit to a 1:1 binding model that accounts for the presence of both free and bound protein:

$$FP = A\frac{([R]_{total} + [P]_{total} + K_D) - \sqrt{([R]_{total} + [P]_{total} + K_D)^2 - 4 \times [R]_{total} \times [P]_{total}}}{2 \times [R]_{total}} + B$$

where A is the amplitude, B is the baseline (minimum FP response), $[R]_{\text{total}}$ is the peptide concentration (in µM), and $[P]_{\text{total}}$ is the protein concentration (in µM).[2] All FP assays were performed in duplicate on two different days; in the rare case when initial results were inconsistent, a third replicate was obtained. Fit parameters for all peptide and protein combinations are provided in Table S2, reported as the average and standard deviation of the parameters from the two replicates for each peptide and protein combination.

**Supplementary References**

(1) O'Neal, L. G.; Drucker, M. N.; Lai, N. K.; Clemente, A. F.; Campbell, A. P.; Way, L. E.; Hong, S.; Holmes, E. E.; Rancic, S. J.; Sawyer, N.; Wang, X.; Thrall, E. S. The *B. Subtilis* Replicative Polymerases Bind the Sliding Clamp with Different Strengths to Tune Their Activity in DNA Replication. *Nucleic Acids Research* **2025**, *53* (14), gkaf721. https://doi.org/10.1093/nar/gkaf721.

(2) Jarmoskaite, I.; AlSadhan, I.; Vaidyanathan, P. P.; Herschlag, D. How to Measure and Evaluate Binding Affinities. *eLife* **2020**, *9*, e57264. https://doi.org/10.7554/eLife.57264.
